# Computational modeling of posteroanterior lumbar traction by an automated massage bed: predicting intervertebral disc stresses and deformation

**DOI:** 10.1101/2022.04.25.489383

**Authors:** Luis Cardoso, Niranjan Khadka, Jacek Dmochowski, Edson Meneses, Youngsoo Jin, Marom Bikson

## Abstract

Spinal traction is a physical intervention that provides constant or intermittent stretching axial force to the lumbar vertebrae to gradually distract spinal tissues into better alignment, reduce intervertebral disc (IVD) pressure, and manage lower back pain (LBP). However, such axial traction may change the normal lordotic curvature, and result in unwanted side effects and/or inefficient reduction of the IVD pressure. An alternative to axial traction has been recently tested, consisting of posteroanterior (PA) traction in supine posture, which was recently shown effective to increase the intervertebral space and lordotic angle using MRI. PA traction aims to maintain the lumbar lordosis curvature throughout the spinal traction therapy while reducing the intradiscal pressure. In this study, we developed finite element simulations of mechanical therapy produced by a commercial thermo-mechanical massage bed capable of spinal PA traction. The stress relief produced on the lumbar discs by the posteroanterior traction system was investigated on human subject models with different BMI (normal, overweight, moderate obese and extreme obese BMI cases). We predict typical traction levels lead to significant distraction stresses in the lumbar discs, thus producing a stress relief by reducing the compression stresses normally experienced by these tissues. Also, the stress relief experienced by the lumbar discs was effective in all BMI models, and it was found maximal in the normal BMI model. These results are consistent with prior observations of therapeutic benefits derived from spinal AP traction.

## Introduction

Lower back pain (LBP) is defined as pain on the posterior aspect of the body from the twelfth ribs to the lower gluteal folds that last for at least one day [1, 2]. LBP affects more than 570 million people globally, equivalent to about 7.5% of the world population in 2017 [3–5]. Back pain may be acute if only last a few days or weeks, and tends to resolve without residual loss of function. In turn, chronic back pain is experienced for 12 weeks or longer, and requires medical treatment or surgery. Most acute back pain cases have a mechanical component, including physical disruption of the spine, muscle, nerves and intervertebral discs (IVD) in the lumbar region of the spine. The mechanical causes of LBP can have different origin, such as: congenital (e.g. spinal bifida, scoliosis, lordosis, kyphosis, etc.), injuries (e.g. tendon/muscle/ligament tears and spasms, traumatic injuries, etc.), degenerative diseases (e.g. intervertebral disc degeneration, spondylosis, arthritis, etc.), nerve and spinal cord problems (e.g. spinal nerve compression, sciatica, stenosis, spondylolisthesis, etc.) and non-spinal factors (e.g. fibromyalgia, tumors, pregnancy, etc.) [1, 2].

Several risk factors have been associated with the development of LBP, among those, age, fitness level, weight (e.g. overweight, obese, etc.) and genetics are the most common. Acute back pain can be treated with medications aimed at reducing pain and inflammation (e.g. analgesics, non-steroidal anti-inflammatory drugs, muscle relaxants, topical pain relief, etc.) thermal therapy (e.g. heat/cold) and gentle exercise and physical stretching. In turn, chronic back pain may require a progressive care approach, starting with early treatments including medication and self-managed thermal therapy. These approaches may be followed by complementary techniques such as acupuncture, transcutaneous electrical nerve stimulation (TENS), physical therapy, spinal manipulation/mobilization, spinal injections, spinal traction and surgery. Mechanical based approaches are generally used for therapeutic purposes including management of pain [6–9], relaxation / autonomic health [10], enhancing local circulation [11–17], and recovery from fatigue [11, 16, 18-20].

Spinal traction is of particular interest because provides constant or intermittent stretching force to the lumbar vertebrae to gradually distract skeletal structures / spinal tissues into better alignment, reduce the intradiscal pressure, and reduce back pain [21–28]. Spinal traction is often used for the treatment of intervertebral disc related problems, comprising herniated discs, sciatica, degenerative disc disease, pinched nerves and LBP [21, 29-31]. Indeed, the mechanism behind mechanical traction of the lumbar spine is believed to include increased blood flow in the tissues, reduced intradiscal pressure, decrease pressure on the nerves and improved stiffness of spinal structures [32].

Different spinal traction approaches have been implemented in the clinic, including: (a) continuous traction, where low weights are applied for extended periods of time; (b) sustained traction, when heavier weights are applied steadily for short periods of time; (c) intermittent mechanical traction, similar to sustained traction, but uses a mechanical system to control the traction force at preset intervals; (d) manual traction, is delivered by the therapist’s hand, sometimes using a belt to pull on the patient’s legs; (e) autotraction, in which the patient pulls or pushes him/herself using a specially designed table that can be tilted or rotated; (f) positional traction, implies placing the subject in appropriate positions using pillows/blocks to create a longitudinal pull on the spinal structures; and (g) gravity lumbar traction, uses a chest harness to support the upper body of the patient, while the lower body weight is used as the traction force. A common feature of these approaches is that they all apply an axial traction force in the craniocaudal direction of the spine. However, it has been shown that axial traction may lead to changes of the normal lordotic curvature [33, 34], decreasing the lumbar lordotic angle and potentially resulting in muscle pain, spasms, joint and interspinous ligament damage, and inefficient reduction of IVD compression [33, 34]. Thus, the clinical usefulness of spinal traction to help with LBP has been unclear [33–35]. An alternative to axial traction has been recently investigated [35–37], based on posteroanterior (PA) traction or the combination of axial traction with PA traction, which aims to maintain the lumbar lordosis curvature throughout the spinal traction therapy [35, 38] and reduce the intradiscal pressure [36]. Posteroanterior traction in supine posture was recently shown using MRI analysis to effectively increase the intervertebral space and lordotic angle [37]. However, the biomechanics behind the spinal PA traction approach has not been fully investigated.

In this study, we develop a state-of-the art simulation of mechanical therapy produced by a commercial thermo-mechanical massage bed (Master V4, CGM MB-1901, CERAGEM Co. Ltd., Cheonan, Korea) capable of posteroanterior spinal traction. We optimized an MRI imaging approach to resolve key back anatomy, developed a 3D model of the relevant tissues (including skin, subcutaneous fat, soft tissue, muscles, intervertebral disc, vertebrae, epidural fat, cerebrospinal fluid, and spinal cord), represented the applied PA traction under static conditions, and used Finite-Element-Method (FEM) to predict relevant effects, including the stresses and strains in the intervertebral discs on the lumbar spine. The mechanical traction is produced in the posteroanterior direction, taking advantage of the spinal lordosis to distract the vertebral bodies, while the subject is laying on the massage bed in supine position, contrary to most spinal traction devices and approaches that use mechanical traction in the craniocaudal direction. The stress relief produced on the lumbar discs by the posteroanterior traction of the device was investigated on human subject models with different BMI (normal, overweight, moderate obese and extreme obese BMI cases). We show significant changes in stress relief in the lumbar discs as a function of the traction level delivered by the device.

## Methods

### MRI scans and Anatomical model

We collected high resolution T2-weighted lumbar spine MRI scans of a healthy male adult (BMI: ~ 25 Kg/m^2^; age: 42 years) using 3T Siemens MAGNETOM Prisma scanner equipped with CP Spine array coil (Siemens Healthineers, PA, USA). Image acquisition parameters were given as: TE: 99 ms; TR: 7040 ms; flip angle: 130°; FOV: 256 mm; SNR: 1; resolution: 256 x 256; slice thickness: 1 mm; and voxel size: 1 x 1x 1 mm. The MR scans were segmented into nine tissue masks: skin, subcutaneous fat, soft tissue, muscles, intervertebral disc, vertebrae, epidural fat, cerebrospinal fluid, and spinal cord. Manual tissue segmentation included using thresholding and morphological filters such as flood fill, dilation, and erosion available in Simpleware ScanIP (Synopsys Inc., CA, USA). Exhaustive review and adjustment ensured state-of-the-art precision of the subject’s spine and surrounding tissue.

The normal BMI model’s subcutaneous fat (thickness: 13 mm) was further artificially dilated to generate models with different BMI values, namely overweight (25 < BMI < 30; thickness: 26 mm), moderate obese (30 < BMI < 40; thickness: 52 mm), and extreme obese (40 < BMI < 65; thickness: 86 mm)[39]. For the dilation procedure, the subcutaneous fat of the normal BMI spine model was first merged with the skin, then dilated isometrically to the aforementioned fat thickness, and later the skin mask was recovered by further dilating the merged mask by the original thickness of the skin (~ 1 mm) to form the new skin mask.

The four BMI models were meshed in Simpleware ScanIP using an adaptive tetrahedral voxel-based meshing algorithm into a finer mesh. The resulting normal BMI model consisted of > 3.36M tetrahedral elements, the overweight BMI model has > 3.43M elements, the moderate obese model comprised >3.46M elements, and the extreme obese model included > 3.68M elements. The four models were later imported into Abaqus (v.2019, Dassault Systems, MA, USA) to computationally solve the finite element method (FEM) model.

### Mechanical actuator

The mechanical actuator of a commercial spinal thermal massage device (Master 4, CGM MB-1901, CERAGEM Co. Ltd., Cheonan, Korea)[38] that provides posteroanterior traction, was investigated using numerical modeling. The mechanical actuator assembly was imported into Abaqus (v.2019, Dassault Systems, MA, USA) for positioning of its parts, creating the rotary linkages (hinges) of several movable components and it was meshed using an adaptive tetrahedral voxel-based meshing algorithm. The mechanical actuator can move horizontally along the craniocaudal direction of the patient laying on supine position on the device bed. The actuator also comprises four semi-cylindrical massage rollers located under the device mat, which move vertically to identify the curvature of the patient’s spine. The actuator then moves to specific locations in the lumbar or cervical regions of the patient’s spine and gradually lift the massage rollers in the posteroanterior direction. The vertical displacement of the actuator is controlled by the traction level (TL) setting of the system. The full range of motion is divided in 9 consecutive TL values, where the vertical displacement of the actuator is increased by approximately 6.9 mm at each traction level, resulting in a maximal vertical displacement of 62 mm at TL9. Fig. 2 shows the mechanical actuator and the massage bed mat at the initial position (Fig. 2A) and at three different vertical position levels (Fig. 2B, 2C and 2D) corresponding to traction level 3, 6 and 9, respectively.

**Figure 1:**
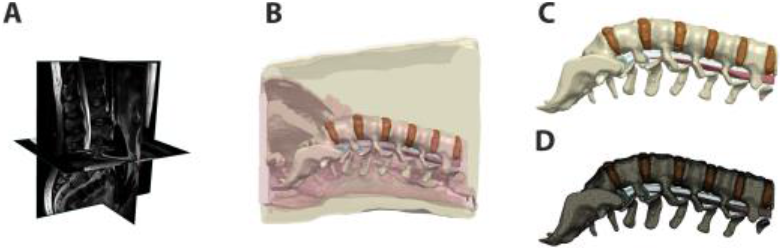
Lower back human model. (**A**) MRI scan of lower back of a normal BMI subject, (**B**) Segmentation of tissues; (**C**) 3D rendering of lumbar discs and spine; (**D**) Finite element mesh of the lumbar discs and spine.

**Figure 2:**
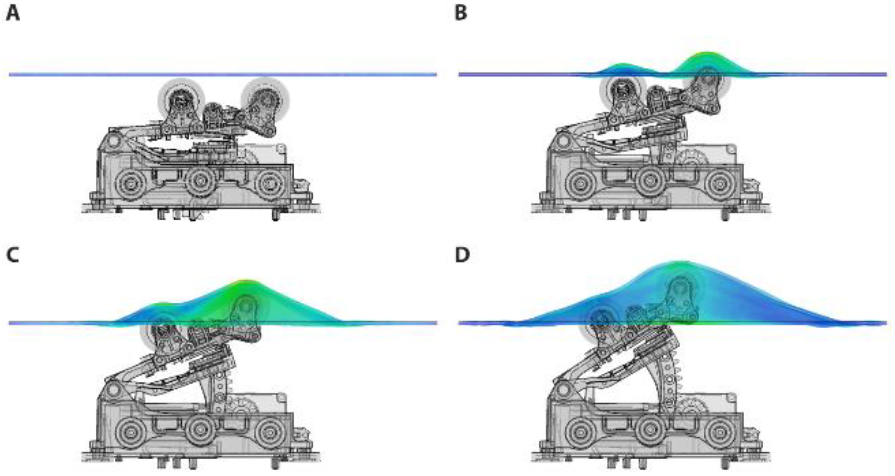
Mechanical actuator and bed mat layer. Vertical position of actuator rollers at different traction levels. Fig 2A shows resting position of actuator, Fig2B represents the vertical displacement at traction level 3 (TL3); Fig 2C shows the position of the actuator and bed mat at TL6; Figure 2D is the maximal height reached by the actuator at TL9, corresponding to ~62mm above resting position.

### Mechanical modeling

An assembly was created in Abaqus between the human subject model and the mechanical actuator. The subject model was positioned directly above the mechanical actuator and the bed mat, with a clearance of 0.1 mm. A contact condition was implemented between the surface of the actuator rollers and the bottom surface bed mat, and a second contact condition was created between the top surface of the bed mat and the bottom (posterior side) of the human subject. The rollers and the top of the bed mat were defined as master surfaces, and in turn, the bottom of the bed mat and the skin in the subject’s back were identified as slave surfaces for the contact condition. The contact between these surfaces was defined as frictionless in the tangential direction and as “hard contact” in the normal direction. A set of nodes were selected on the top and bottom, anterior-aspect of the human subject, and defined as a boundary condition with zero translation in three spatial coordinates, but free to rotate in any direction. Each tissue constituent was assigned linear elastic material properties previously reported in the literature[40–49]: (**1**) skin (E = 160 MPa; ν = 0.49; ρ = 1020 kg m-3), (**2**) muscle (E = 7 MPa; ν = 0.49; ρ = 1100 kg m-3), (**3**) soft tissue (E = 23.5 MPa; ν = 0.49; ρ = 1057 kg m-3), (**4**) vertebrae (E = 17 GPa; ν = 0.30; ρ = 1,800 kg m-3), (**5**) intervertebral disc (E = 17 MPa; ν = 0.49; ρ = 1100 kg m-3), (**6**) subcutaneous fat/epidural fat: (E = 3 MPa; ν = 0.49; ρ = 920 kg m-3), (**7**) CSF: (K = 2.25 GPa; ν = 0.499; ρ = 1000 kg m-3), (**8**) spinal cord/dura mater: (E = 10 MPa; ν = 0.49; ρ = 1057 kg m-3). A dynamic explicit formulation was used to solve for the deformation, stresses and strains produced in the human subject model by the displacement of the mechanical actuator.

## Results

### Mechanical actuation on normal BMI model

A mid-sagittal view of the effect produced by mechanical actuation on the tissues of the normal BMI subject model at traction level 9 is shown in Fig. 3. The 3D stresses are depicted in Fig. 3A and the strains in Fig. 3B. This result shows that the largest stresses and deformation occurs at the center of the lumbar spine, at the level of the L2-L4 vertebral bodies. The stress plot shows von Mises stresses, demonstrating the heterogeneity of stresses in the lower back under the action of the mechanical traction. As expected, the highest stresses and smallest deformations occur at the vertebral bodies, since calcified tissues have the highest elastic modulus associated with them. The internal stresses in the intervertebral discs increase as a function of traction level. These stresses counteract the compression stress level in the disc prior to the mechanical traction.

**Figure 3.**
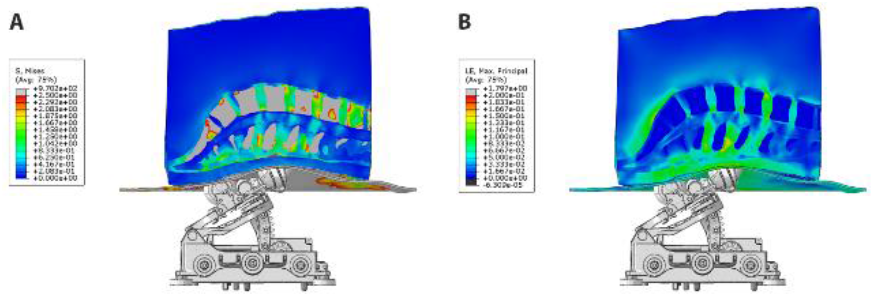
Actuator and model assembly. Fig. 3A shows the stresses and Fig. 3B depicts the strains produced by posteroanterior traction produced by the system on the tissues of a normal BMI subject. The front rollers produce the highest deformation on the skin and fatty tissues layer, as well as maximal traction directly under the location of the vertebral bodies L2 to L4.

### Comparison of 3D stress map on different BMI models

The 3D stress maps on different BMI models: normal, overweight, moderate obese and extreme obese (column panels) are compared at traction levels 5, 7 and 9 (row panels) in Fig. 4. In these panels, we observe that the highest stresses in the different tissues of the model occur in the normal BMI model at the highest traction level. While the effect of mechanical traction is also apparent in the overweight, moderate obese and severe obese models, the relative intensity of stresses decreases as a function of BMI. Analysis of the 3D stress maps of the lumbar discs and spine of different BMI models demonstrate that the effect of mechanical traction is evident in all models. The normal BMI is the model with the highest stress relief in the lumbar discs, and the intensity of stress relief is inversely proportional to the BMI in the model; still the mechanical traction of the system is present even in the severe obese BMI model.

**Figure 4.**
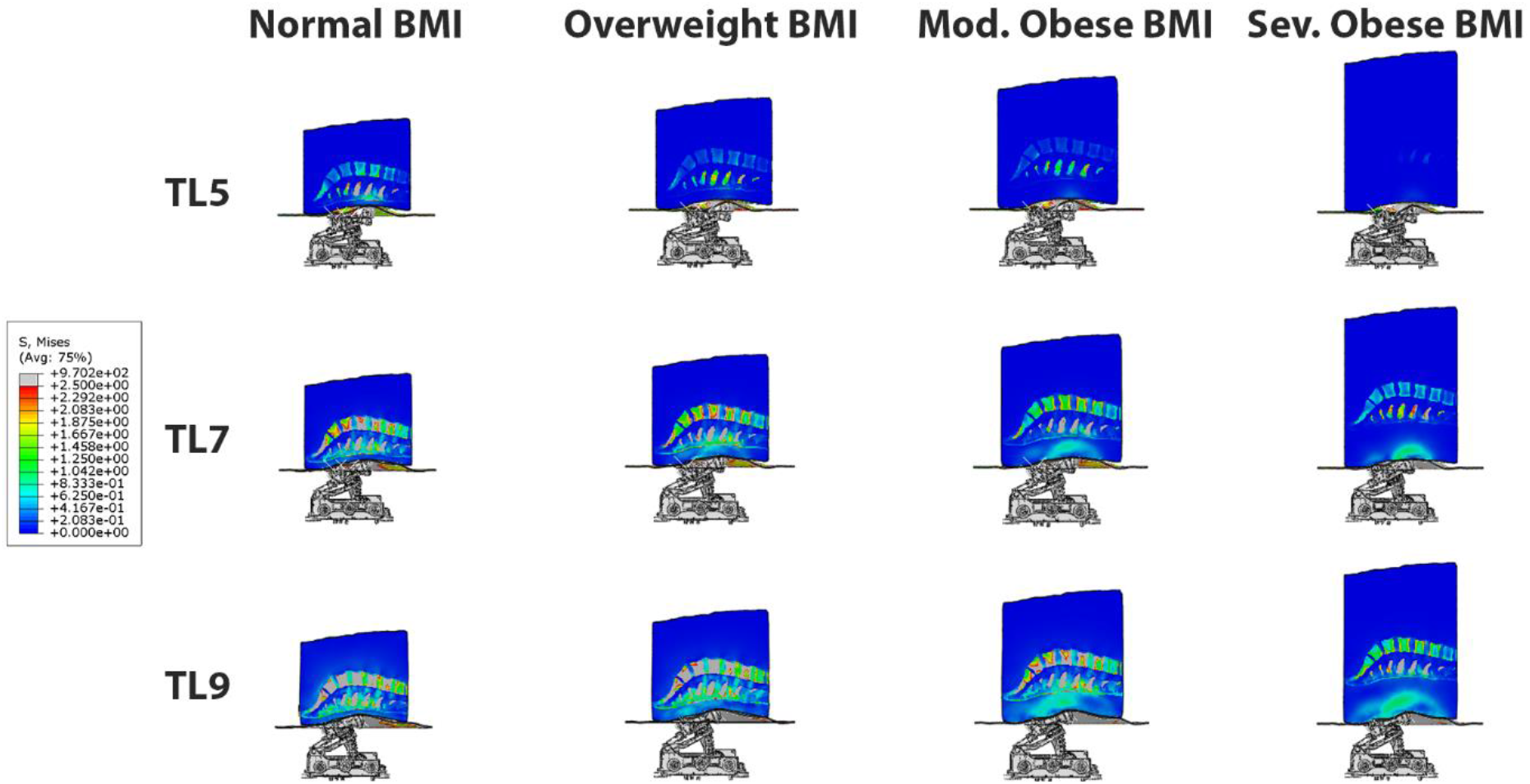
3D Stress map on different BMI models: normal, overweight, moderate obese and severe obese (column panels) at traction levels 5, 7 and 9 (row panels). The highest stresses occur in the normal BMI model at the highest traction level. The effect of posteroanterior traction is inversely proportional to BMI, and it is evident in all BMI models.

### Analysis of 3D Stresses on Different BMI Models as a function of traction level

A quantitative comparison of 3D internal stresses produced on different BMI models: normal, overweight, moderate obese and severe obese as a function of traction level (TL 1-9) is presented in Fig. 5. These curves were obtained by averaging the stresses in all the nodes of each of the six lumbar discs (approx. 20,000 nodes per disc) as a function of mechanical traction level. The stresses in the models remain close to zero value for TL4 and lower, since the actuator rollers start contact with the lower back at this level. The internal stresses developed in the lumbar discs exhibited a range of variability between 0.075 and 1.7 MPa, depending on the BMI and traction level (TL5-TL9). The normal BMI model exhibit the lowest stresses in the T12-L1 disc (~0.35MPa), followed by the L5-S disc (~0.8MPa). A similar level of maximal stresses was found in the L1-L2 and L4-L5 discs (~1.2MPa), a little higher in the L3-L4 disc (~1.5MPa) and the highest stresses were observed in the L2-L3 disc (~1.8MPa). The apparent increase on stresses as a function of traction level at the different lumbar discs is nonlinear, possibly due to the complex / inhomogeneous deformation of the several tissues considered in the model and the morphology / curvature of the spine, even though the tissues mechanical properties were defined using linear elastic models. The presence of fat tissue in the posterior aspect of the lower back reduces the magnitude of the stresses seen in the intervertebral discs. The overweight BME model exhibit similar trends in disc stresses when compared to the normal BMI model, but with stress magnitude reduced by 10-20% approximately. An equivalent behavior is recognized for the moderate and severe obese BMI models, but with a stress magnitude reduced by about 50% and 75%, respectively. This quantitative result represents a cause-effect relationship between the mechanical traction of the massage bed and the stress relief in each lumbar disc. In general, the L2-L3 disc was the one with the highest stresses, and the T12-L1 disc had the lowest stresses. These curves confirmed the qualitative observation in Fig. 4 indicating that the internal stresses in the disc are maximal in the normal BMI model and decrease as the BMI increases.

**Figure 5.**
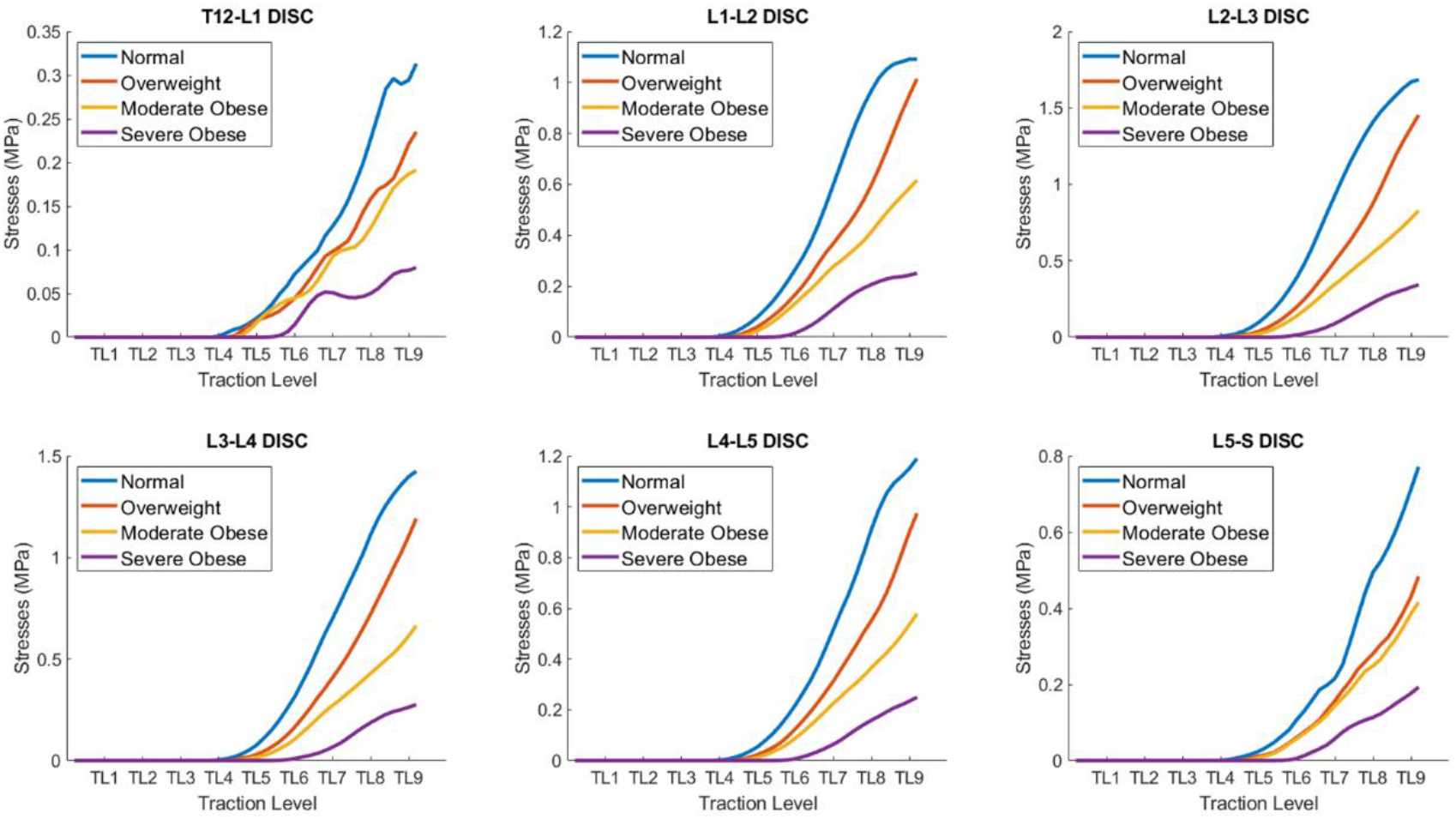
Quantitative comparison of average stresses on different BMI models: normal, overweight, moderate obese and extreme obese as a function of traction level (TL 1-9) in the lumbar discs. The internal stresses developed in the lumbar discs exhibit a range of variability between 0.075 and 1.7 MPa, depending on the BMI and traction level. The internal stresses in the disc are maximal in the normal BMI model and decrease as the BMI increases.

### Comparison of 3D strain map on different BMI models

The behavior of mechanical strains is equivalent to stresses in all tissues and models, as a function of traction level and BMI. Fig. 6 shows 3D strain maps on different BMI models: normal, overweight, moderate obese and extreme obese (column panels) at traction levels 5, 7 and 9 (row panels). Similar to the stresses result, the strains in all models for the TL4 or lower are practically zero, since the actuator rollers touches the lower back tissues when the actuator moves from TL4 to TL5. At TL5 we can readily observe deformation of the lower back. In these panels, we can observe similar trends shown in Fig. 4, where the maximal strains in tissues are presented in the normal BMI model at the highest traction level. We can also observe that the maximal strains in the intervertebral discs are presented in the normal BMI model at the highest traction level. However, the fat and soft tissues in the moderate and extreme obese models deform the most and thus reduce the strains observed in the intervertebral discs in high BMI models.

**Figure 6.**
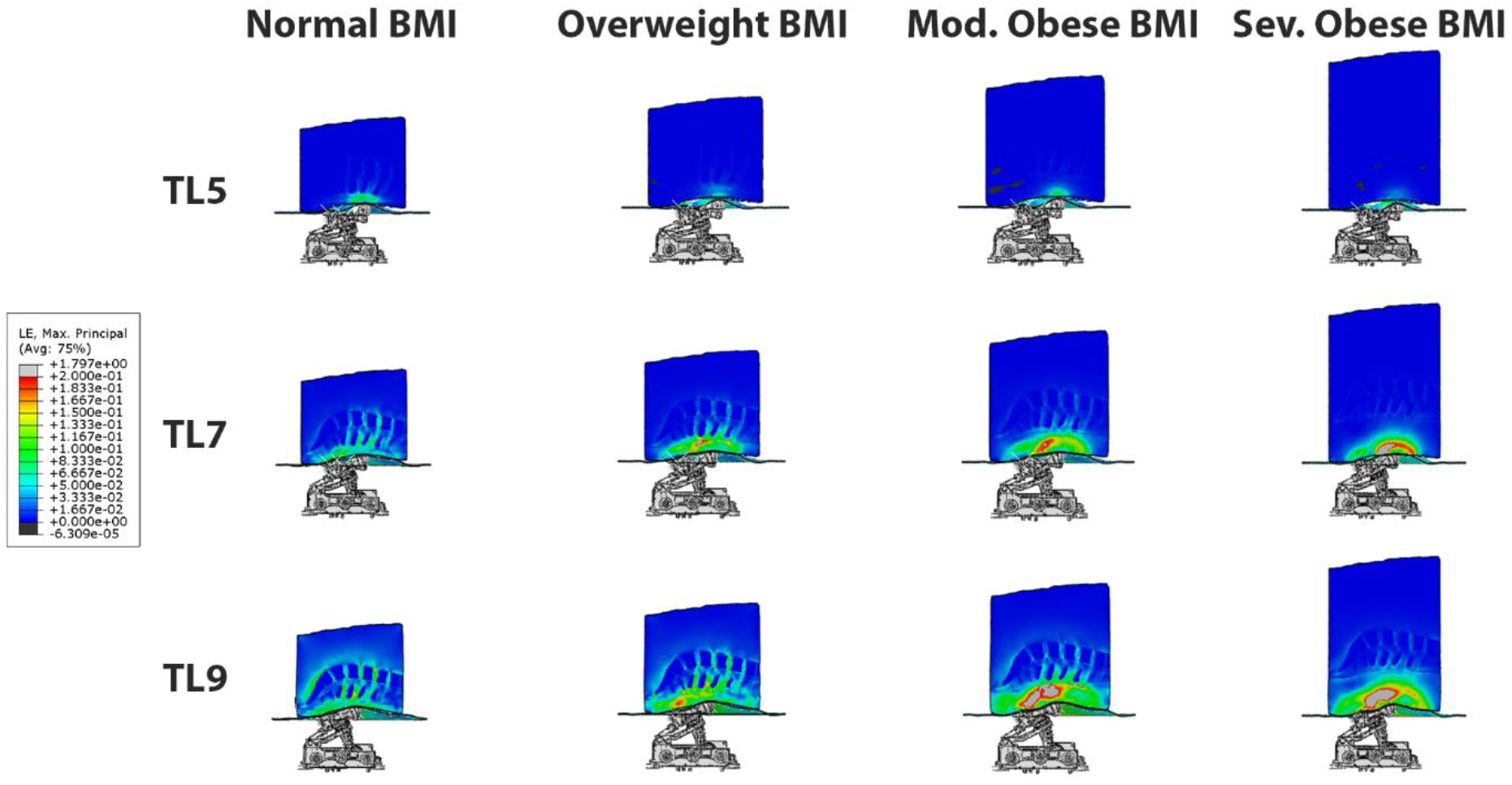
3D strain maps on different BMI models: normal, overweight, moderate obese and extreme obese (column panels) at traction levels 5, 7 and 9 (row panels). The maximal strains in the intervertebral discs are presented in the normal BMI model at the highest traction level. However, the fat and soft tissues in the moderate and extreme obese models deform the most and shield the intervertebral discs from the strains.

### Analysis of 3D Strains on Different BMI Models as a function of traction level

The quantitative comparison of 3D strains produced on different BMI models as a function of traction level is shown in Fig. 7. The approach to quantify the average strain level in the intervertebral discs is similar to the one described earlier for measurement of internal stresses. Overall, we can observe that at TL 9, the average strains range from 0.004 to 0.1 for the severe obese BMI and the normal BMI models, respectively, but vary depending across the different discs. In particular, these results indicate that the L2-L3 disc is the one with the highest deformation (up to 0.1 strain), and the T12-L1 disc has the lowest strains. The curves for the different discs are indeed similar to those obtained for the stresses, and the trends are thus similar too. The maximal strains in the T12-L1 disc are the smallest among all the lumbar discs (~0.018), followed by the L5-S disc (~0.045). The maximal strains are larger in the L1-L2 disc (~0.06), the L4-L5 disc (~0.07) and the L3-L4 disc (~0.08), and are maximal in the L2-L3 disc, reaching a strain of about 0.1. Similar to the stresses, the strains in the overweight BMI model is 10-20% lower than in the normal BMI model. The moderate obese BMI model has about 50% lower strains when compared to the normal BMI model, and the severe obese BMI model shows ~75% lower strains than the normal model. Thus, these curves also confirm that the internal strains in the disc are the highest in the normal BMI model and decrease as the BMI increases.

**Figure 7.**
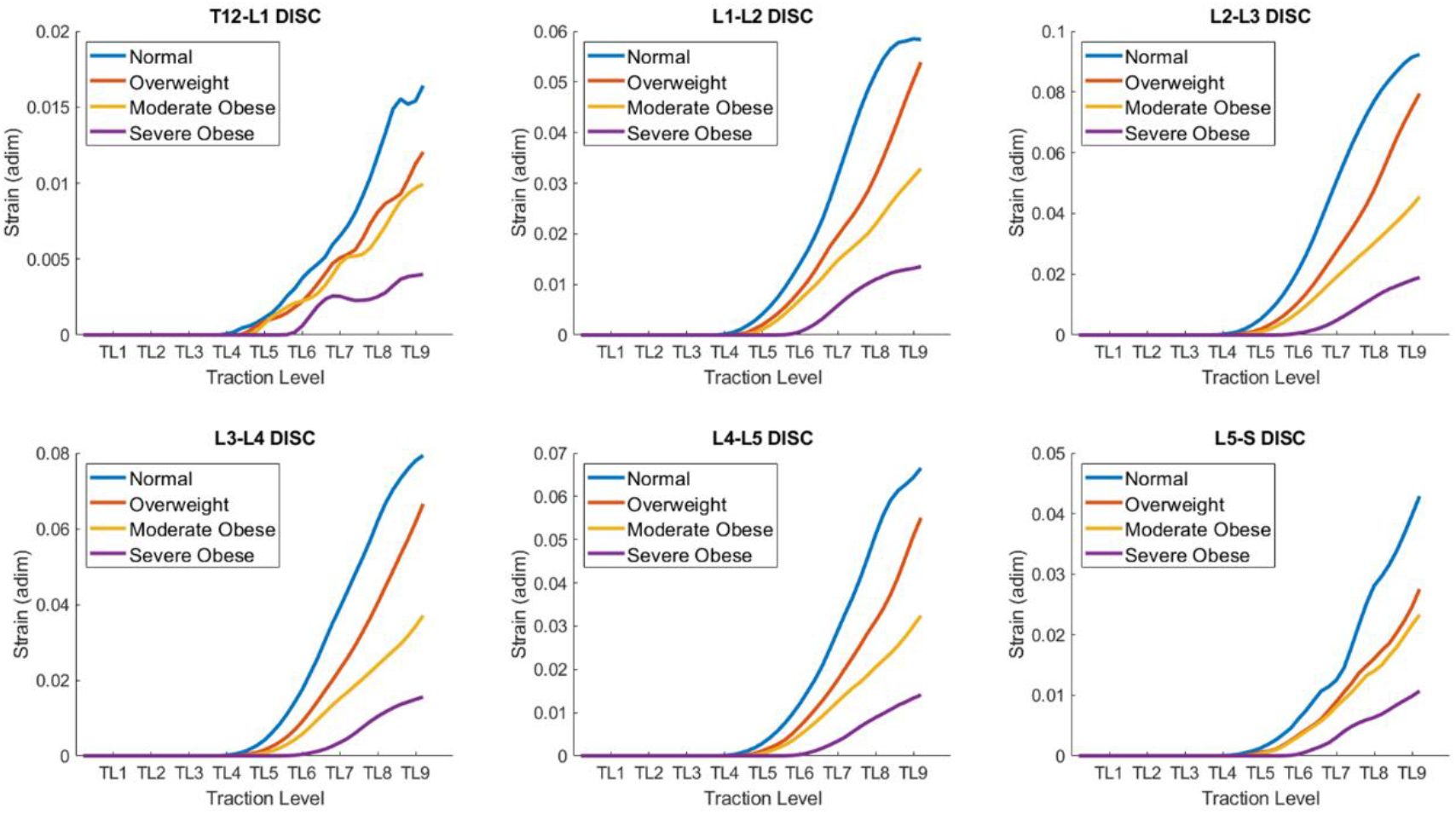
Quantitative comparison of Strains on different BMI models: normal, overweight, moderate obese and extreme obese as a function of traction level (TL 1-9) in the lumbar discs. The strains developed in the lumbar discs exhibit a range of variability between 0.005 and 0.1, depending on the BMI and traction level. The strains in the disc are maximal in the normal BMI model and decrease as the BMI increases.

## Discussion

Prolonged massage therapy has an extensive history in wellness and medical treatment but recent advances in automatic massage beds suggest new potency for self-managed care. Evidently, the outcomes of mechanical massage will depend on the technical features of the mechanical actuators, the type of mechanical traction, as well as the individual’s anatomy and physiology. In this study we performed a quantitative computational analysis of the internal stresses and strains produced in the lumbar intervertebral discs by a system that delivers posteroanterior traction of the spine. We tested the effect of different traction levels in four models with different BMI, and demonstrated the effectiveness of the approach to produce distraction of the lumbar discs, in agreement with changes of lordotic angle reported in human subjects using MRI [37]. Our study provides supporting evidence on the distraction effect of posteroanterior traction for all lumbar discs in different BMI subject models.

The predictions here also provide a direct substrate for observations of changes in intervertebral space [37] and the relief of internal compressive stresses via the induction of distraction stresses that counterbalance the compressive ones, which in turn can decrease lower back pain and disc compression states that precedes disc degeneration. The stress and strain levels predicted by the models in the intervertebral discs are below the reported thresholds for damage of the disc (ultimate stress = 2.94±1.05MPa and ultimate tensile strains = 21.3±2.1%) [47], consistent with the established safety of the commercial automatic massage bed system. It is further shown that the mechanical traction delivered by this system is effective in a broad range of BMI subjects, and it is directly proportional to the set traction level of the therapy. In addition, tensile stresses produced by this device in the disc tissue can explain clinical observations using the same automatic message devices modeled here on immune status [6], autonomic function [10], inflammation[50, 51], pain [6-9, 52, 53], intervertebral space changes [37], and activate antioxidant enzymes [7].

Computational models of medical devices relate device features (set by the operator and outside the body) with resulting changes in tissue properties – which in turn govern therapeutic actions. Therefore, the mechanical effects predicted here have direct implications on understanding (and further optimizing) results from clinical trials using the same device. Evidently, the predictions reported here are specific to the mechanical traction modeled, for example segments of maximal strains (Fig. 7) and stresses (Fig. 5) correspond to front roller position (Fig. 3). Our result support general inferences regarding the mechanisms of the mechanical massage technology simulated.

There are several valuable next steps to further this study, which will include 1) directly verify model predictions by physiologic measurements; and 2) suggesting improved protocols whose clinical benefits can then be validated. The present study considered only posteroanterior traction, and future modeling efforts should integrate stress/heating multi-physics in a dynamic model–including considering synergists actions on tissue and clinical outcomes. We did not consider the theoretical impact of deviating from isotropic and linear elastic tissue material properties. Also, the craniocaudal direction positioning of the actuator relative to the lumbar spine undoubtedly play an important role in the produced traction, and other positions should be investigated.

## Author contributions

LC designed the research, performed the research, and wrote the manuscript. NK designed the research, performed the research, and wrote the manuscript. JD designed the research. EM performed the research. YS designed the research and wrote the manuscript. MB designed the research and wrote the manuscript.

